# Premature MeCP2 Expression Disturbs the Subtype Specification of Midbrain Dopamine Neurons

**DOI:** 10.1101/2021.11.12.468365

**Authors:** Xi-Biao He, Fang Guo, Kexuan Li, Jiaqing Yan, Sang-Hun Lee

## Abstract

Midbrain dopamine (DA) neurons are associated with locomotor and psychiatric disorders. DA subtype is specified in ancestral neural precursor cells (NPCs) and maintained throughout neuronal differentiation. Here we show that endogenous expression of MeCP2 coincides with DA subtype specification in mouse mesencephalon, and premature expression of MeCP2 prevents in vitro cultured NPCs from acquiring DA subtype through interfering NURR1 transactivation of DA phenotype genes. By contrast, MeCP2 overexpression does not disturb DA subtype in DA neurons. By analyzing the dynamic change of DNA methylation along DA neuronal differentiation at the promoter of DA phenotype gene tyrosine hydroxylase (*Th*), we show that *Th* expression is determined by TET1-mediated de-methylation of NURR1 binding sites within *Th* promoter. Chromatin immunoprecipitation assays demonstrate that MeCP2 dominates the DNA binding of the corresponding sites thereby blocking TET1 function in DA NPCs, whereas TET1-mediated de-methylation prevents excessive MeCP2 binding in DA neurons. The significance of temporal DNA methylation status is further confirmed by targeted methylation/demethylation experiments showing that targeted de-methylation in DA NPCs protects DA subtype specification from MeCP2 overexpression, whereas targeted methylation disturbs subtype maintenance in MeCP2-overexpressed DA neurons. These findings suggest the appropriate timing of MeCP2 expression as a novel determining factor for guiding NPCs into DA lineage.

## Introduction

Neurogenesis is characterized by generic neuronal differentiation and neurotransmitter subtype specification of neural precursor cells (NPCs), two developmental programs contributing to the molecular, cellular and functional specificity of different neurons in CNS. These two programs are under precise spatio-temporal control of a variety of extrinsic cues and intrinsic transcription factors and epigenetic modifications. Some transcription factors contribute to each program independently, whereas others are involved in both. For instance, in the neurogenesis of midbrain dopamine (DA) neurons, mutation of nuclear orphan receptor gene *Nurr1* impairs the DA subtype specification but not the general neuronal differentiation of DA NPCs (Zetterström, 1997), whereas lack of proneural gene *Ngn2* results in failed acquisition of DA subtype and general neuronal differentiation (Andersson, 2006). Understanding how neuronal subtype specification and general neuronal differentiation are coordinated to reach a complete neurogenesis provides the fundamental basis for the development of stem cell therapy which aims at generating specific neuronal cells with full functionality from embryonic stem cells and induced pluripotent stem cells.

The subtype specification of midbrain DA neurons is transcriptionally characterized by constitutive expression of DA phenotype genes such as *Th* encoding the DA synthesis rate limiting enzyme tyrosine hydroxylase and *Aadc* encoding aromatic L-amino acid decarboxylase, both of which are responsible for the conversion of L-tyrosine into DA. *Th* is also one of the earliest expressed DA phenotype genes that is maintained throughout DA neuron lifespan. The subtype specification of midbrain DA neurons marks the fate commitment of NPCs into DA lineage in embryonic brains, and the maintenance of DA phenotype is essential for mature DA neuronal functions, loss of which in adult midbrains is associated with movement and emotional disorders such as Parkinson’s disease and schizophrenia (Arenas et al., 2015; Goridis and Rohrer, 2002; Hynes and Rosenthal, 1999). Nuclear receptor related 1 protein (NURR1) is a master regulator for the subtype specification of midbrain DA neurons, expression of which in mouse is initiated in DA NPCs residing in the ventral mesencephalon at embryonic day E10.5 (Ang, 2006). Continuous activation of *Nurr1* is implicated in the subtype specification as well as the maintenance of midbrain DA neurons. Genetic ablation of *Nurr1* in transgenic mice resulted in absence of DA phenotype (Saucedo-Cardenas et al., 1998; Zetterström, 1997). Conditional ablation of *Nurr1* caused loss of DA phenotype in maturing DA neurons (Kadkhodaei et al., 2009). By contrast, ectopic expression of *Nurr1* is able to force the expression of these markers in non-DA cells(Kim et al., 2003a; Shim et al., 2007; Wagner et al., 1999). A large number of studies including ours have demonstrated that NURR1 facilitates DA subtype specification through direct transactivation of the promoter activity of DA phenotype genes including *Th* and *Aadc* (He et al., 2011, 2015; Kim et al., 2003b; Park et al., 2008; Yi et al., 2014). Based on this well-elucidated mechanism, various transcription factors and epigenetic factors have been identified to cooperate with NURR1 in DA subtype specification (van Heesbeen et al., 2013), including DNA hydroxymethylase ten-eleven translocation 1 (TET1) and TET1-induced DNA de-methylation (He et al., 2015). However, the implication of DNA methylation and DNA methylation-related molecules in NURR1-mediated DA subtype specification remains to be elucidated.

X-linked gene *Mecp2-*encoded Methyl-CpG binding protein 2 (MeCP2) recognizes and binds to methylated and unmethylated CG dinucleotide to modulate chromatin structure and gene expression. Genetic loss or gain of *Mecp2* is associated with neurological and developmental disorders Rett syndrome and *Mecp2* duplication syndrome, respectively. In developing brain, MeCP2 expression highly correlates with neurogenesis and is involved in general neuronal differentiation and synapse plasticity (Cassel et al., 2004; Chen, 2003, 2003; Jung et al., 2003; Matarazzo et al., 2004; Tate et al., 1996; Tsujimura et al., 2009). Evidence exist that have suggested the association of MeCP2 with DA-related disorders. For instance, deficit of MeCP2 expression in the brain is associated with DA metabolites abnormality in patients and animal models of Rett syndrome (Gantz et al., 2011; Ide et al., 2005; Kao et al., 2015; Panayotis et al., 2011b, 2011a; Samaco et al., 2009; Zhao et al., 2013). Underlying mechanisms include autonomous deficits of key DA synthesizing enzymes including TH in DA neurons of substantia nigra (SN) and striatum, and non-autonomous effects derived from serotonin neurons. *Mecp2* duplication syndrome, a neurodevelopmental disorder shares many similar behavioral and molecular deficits with Rett syndrome, including abnormal DA levels and motor function (Na et al., 2012; Peters et al., 2013). Although the roles of MeCP2 for general neuronal maturation and functions have been widely revealed, little is known about the contribution of MeCP2 to the subtype specification and maintenance of DA neurons.

We previously have shown that the dissociation of MeCP2 from promoters of DA phenotype genes is required to facilitate NPCs into DA lineage (He et al., 2011). This finding was extended in this study by inducing MeCP2 expression before or after DA subtype specification to elucidate consequences of abnormal timing and dose of MeCP2 expression to the subtype specification and maintenance of midbrain DA neurons.

## Results

### NURR1 precedes MeCP2 expression in DA neurogenesis

The initiation of NURR1 expression in the ventricular zone of mouse ventral mesencephalon at E10.5 marks the identity of undifferentiated DA NPCs and start of DA subtype specification, followed by ventral-lateral migration of these cells and immediate expression of DA phenotype gene *Th* in the intermediate and mantle zones at E12.5. To understand the association of MeCP2 and development of DA neuronal subtype, the expression pattern of NURR1 and MeCP2 was examined by immunofluorescence analysis in ventral regions of E10.5, E12.5 and postnatal day 28 (P28) mouse midbrains, where reside DA NPCs before subtype specification, immature DA neurons after subtype specification and mature DA neurons after differentiation, respectively (Ang, 2006). At E10.5, NURR1 was already expressed in the ventricular zone but not co-expressed with TH, confirming that these cells were DA NPCs not yet undergoing DA subtype specification. Notably, these NURR1+ DA NPCs were negative for MeCP2 (Fig. 1A). At E12.5, however, MeCP2 started to get expressed by NURR1+ cells around the midline of intermediate zone and in the mantle zone, in which TH was also expressed. These results demonstrated a sequential expression of NURR1 and MeCP2 across the period of DA subtype specification, and suggested a coincidence of MeCP2 expression and onset of DA subtype specification. Consistent with this, MeCP2 was expressed in all the TH+Nurr1+ mature DA neurons in ventral tegmental area (VTA) and SN at P28. These in situ expression analyses demonstrated a concurrent onset of MeCP2 expression and DA subtype specification, both of which were preceded by NURR1 expression for about one embryonic day in mouse mesencephalon (depicted in Fig. 1B).

**Figure 1.**
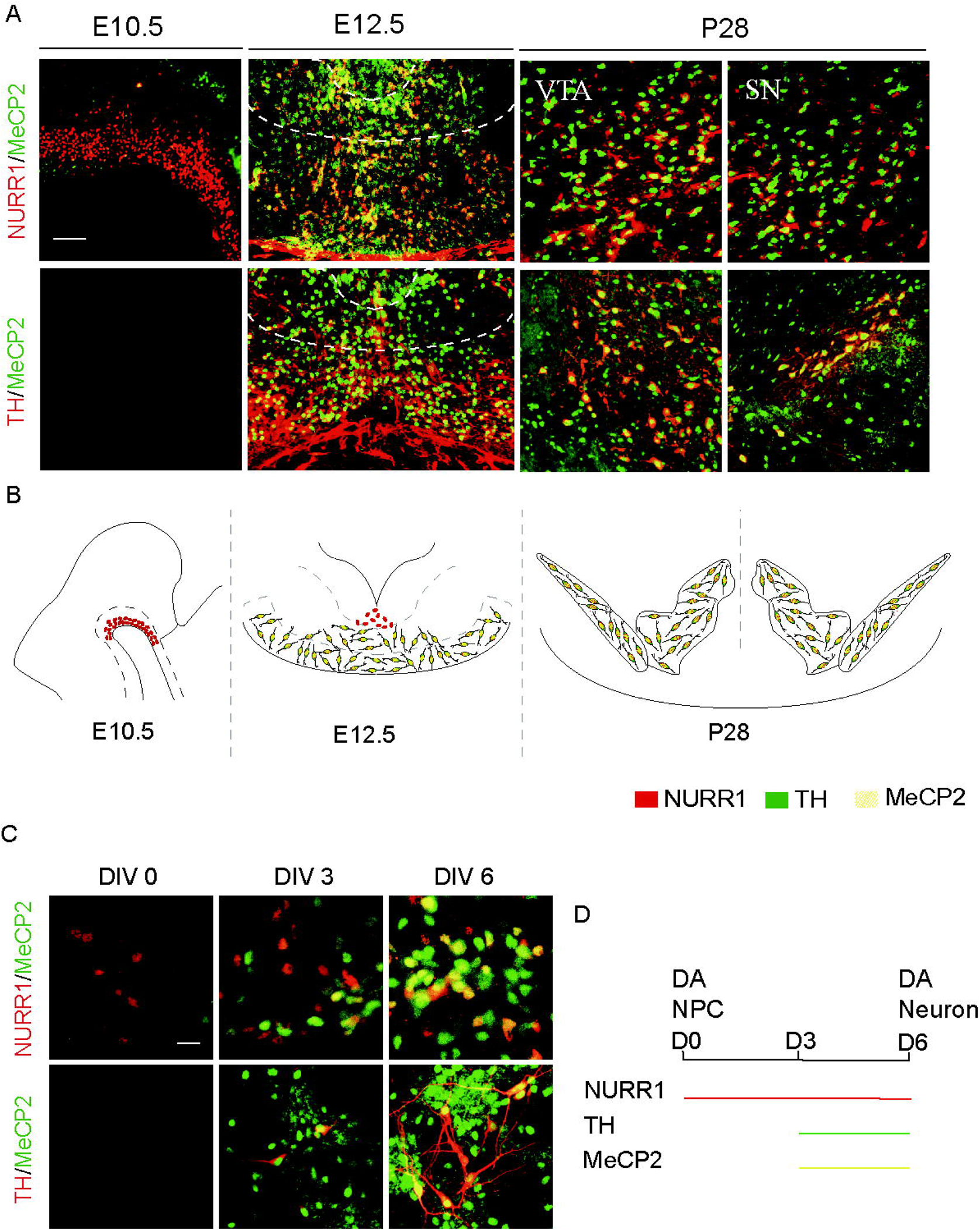
Expression patterns of NURR1, MeCP2 and TH in developing and adult mouse midbrains and cell model of embryonic mouse midbrain dopamine (DA) neuron differentiation. (A) Sagittal (Embryonic day (E) 10.5) and coronal (E12.5 and Postnatal day (P) 28) views of MeCP2-positive cells co-express NURR1 or TH in mouse ventral midbrains. TH is not detected at E10.5. Both ventral tegmental area (VTA) and substantia nigra (SN) of midbrain is shown at P28. Scale bar represents 100 μm. A summary of the expression patterns in vivo is depicted in (B). (C) Cultured DA neural precursor cells (NPCs) were allowed for 6 days of DA neuronal differentiation in vitro (DIV). DIV 0, 3 and 6 were selected for immunofluorescent labeling of cells expressing NURR1/MeCP2 and TH/MeCP2. Scale bar represents 20 μm. A summary of the expression patterns in vitro is depicted in (D).

In our lab, mitotic DA NPCs extracted from mouse ventral mesencephalon at E10.5 can be expanded and induced to differentiate into midbrain DA neurons within 6 days of neuronal differentiation in vitro (DIV), serving as a well-defined cell model to study the mechanisms underlying subtype specification and neuronal differentiation of midbrain DA neurons (He et al., 2011, 2015). This model was recruited to confirm the sequential expression of NURR1 and MeCP2 observed in embryonic development. At DIV0, NURR1+TH- DA NPCs were not expressing MeCP2, whereas NURR1+TH+ DA neurons at DIV3 and DIV6 were expressing MeCP2 (Fig. 1C). Therefore, the DA neuron differentiation cell culture well recapitulated the midbrain DA neuron development in vivo (depicted in Fig. 1D). These results collectively demonstrate a dynamic appearance of endogenous MeCP2 throughout the neuronal differentiation of DA NPCs towards DA neurons. In particular, in DA NPCs, MeCP2 was not expressed until DA subtype was specified, suggesting a hypothetical necessity of restricting MeCP2 expression in DA NPCs for the proper specification of DA subtype.

### Premature MeCP2 expression abolishes DA subtype specification

To test this hypothesis, a lentivirus carrying sequence encoding a tetracycline-inducible MeCP2 was transduced into cultured mitotic NPCs before DA neuronal differentiation. Time- and dose-specific MeCP2 overexpression was induced by addition of 0.1-2 μM doxycycline (Dx) at first three days of DIV, when either endogenous MeCP2 or DA phenotype marker TH was not expressed yet. In parallel, same Dx treatment was applied at last three days of DIV to induce an overdose of MeCP2 after DA subtype specification. At DIV6, the yield of TH+ cells representing differentiated DA neurons was measured (Fig. 2A). We found that MeCP2 overexpression in early period of differentiation remarkably reduced the terminal yield of TH+ cells (Fig. 2B). Interestingly, 0.1 μM Dx resulted in comparable yield reduction of TH+ cells as 2 μM Dx, suggesting no additional influence from dosage of ectopic MeCP2. By contrast, the yield of TH+ cells was not affected by MeCP2 overexpression in late period of differentiation. Molecular analysis demonstrated that the mRNA levels of *Th* and *Aadc* were decreased only when MeCP2 was overexpressed in early but not late period of differentiation (Fig. 2C), suggesting that the subtype specification in DA NPCs was transcriptionally regulated by MeCP2. Collectively, these results demonstrate differential consequences of MeCP2 overexpression to the subtype specification and maintenance of DA neurons.

**Figure 2.**
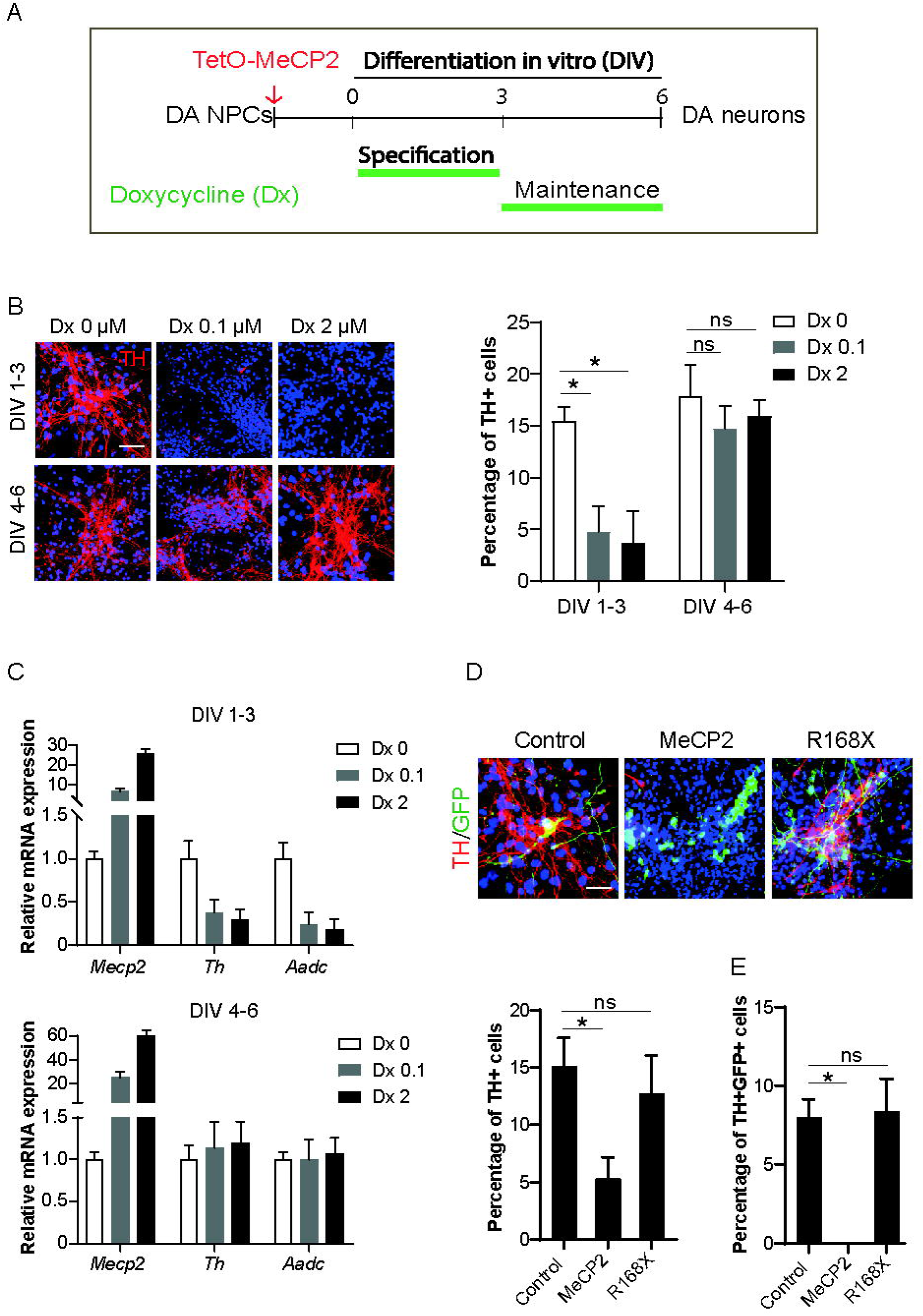
Overexpressing MeCP2 in different periods of dopamine (DA) neuron differentiation. (A) Schematic overview depicting time- and dose-dependent overexpression of MeCP2 by FUW-TetO-mMeCP2 lentivirus within 6 days of DA neuronal differentiation in vitro (DIV). Two concentrations of Doxycycline (Dx) were treated from early (DIV 1-3) or late (DIV 4-6) period of differentiation, in which DA subtype was specified and maintained, respectively. (B) Representative images and quantification of percentage of cells expressing DA subtype marker tyrosine hydroxylase (TH) after 6 days of differentiation. (C) Transcriptional changes of DA phenotype genes *Th* and *Aadc* in response to MeCP2 overexpression. 2 μM Dx was added within DIV 1-3 or DIV 4-6. Analysis was performed DIV 6. One representative data from three independent experiments is shown. Data are expressed as mean ± S.D. (D) Representative images and quantification of percentage of cells expressing TH showing that overexpression of MeCP2 and its transcription repression domain mutation R168X in cultured DA neural precursor cells (NPCs) have different impact on DA subtype specification. Ectopic genes were delivered by lentiviruses carrying ZsGreen1 fluorescent protein (GFP). (E) Quantification of cells double positive for TH and GFP. Note that no double positive cells in MeCP2-overxpressed group, indicating no cells overexpressing MeCP2 acquired DA phenotype after differentiation. Scale bar represents 50 μm. Cell numbers were counted in 10 random areas of each culture coverslip. Data represent mean ± S.E.M. n=3 independent culture. * *P* < 0.05, ns, not significant; one-way ANOVA with Tukey’s post hoc analysis.

Protein structure studies have identified two functional domains in MeCP2, namely DNA binding domain (DBD) and transcriptional repressing domain (TRD). To further dissect the domain functionality of MeCP2 in DA subtype specification, DA NPCs were infected with lentiviruses carrying sequences encoding full-length MeCP2 or R168X, a truncated mutation containing intact DBD but not TRD. A sequence encoding ZsGreen1 fluorescent protein (GFP) was incorporated to label the infected cells. We found that overexpression of MeCP2 from DIV0 significantly reduced the yield of TH+ cells at DIV6, verifying the results observed from Dx-induced MeCP2 overexpression. However, overexpression of R168X did not alter the yield of TH+ cells (Fig. 2D), indicating that the TRD of MeCP2 is responsible for the disturbance of DA subtype specification in DA NPCs. Notably, TH and GFP double positive cells were found in cells infected by empty vector or R168X, but not in cells infected by MeCP2 (Fig. 2E), indicating that MeCP2 disturbed DA subtype specification in a cell-autonomous manner.

### MeCP2 impedes NURR1 transactivation of DA phenotype genes in DA NPCs

NURR1 overexpression is sufficient to induce expression of DA phenotype genes especially *Th* even in non-DA lineage cells such as rodent embryonic cortical NPCs. We recruited this system to gain a direct insight into the association of MeCP2 and NURR1 in DA subtype specification. As the TH-inducing effect has been reported much more effective in NPCs derived from rat than mouse (Lee et al., 2009; Park et al., 2008), rat embryonic cortical NPCs were co-infected with lentivirus encoding NURR1 and that encoding MeCP2 or R168X. About 50% of cells infected with NURR1 and control viruses started to express TH within 3 days, confirming the efficiency of NURR1-induced TH expression. MeCP2 overexpression greatly reduced this population to about 2% (Fig. 3A-B). Co-labeling of TH and GFP showed that the majority of cells overexpressing MeCP2 were negative for TH (Fig. 3C), consistently indicating a cell-autonomous effect of MeCP2. Overexpression of R168X also did not affect the yield of TH+ cells. Real-time PCR analysis demonstrated a transcriptional repression of *Aadc* as *Th* by MeCP2 but not R168X, a result similar to the overexpression outcome in DA NPCs (Fig. 3D). Given that NPCs from embryonic cortices do not possess the majority of molecular machinery for DA neuron development, it appears safe to conclude that ectopic MeCP2 targets NURR1-mediated molecular machinery to inhibit DA phenotype gene expression.

**Figure 3.**
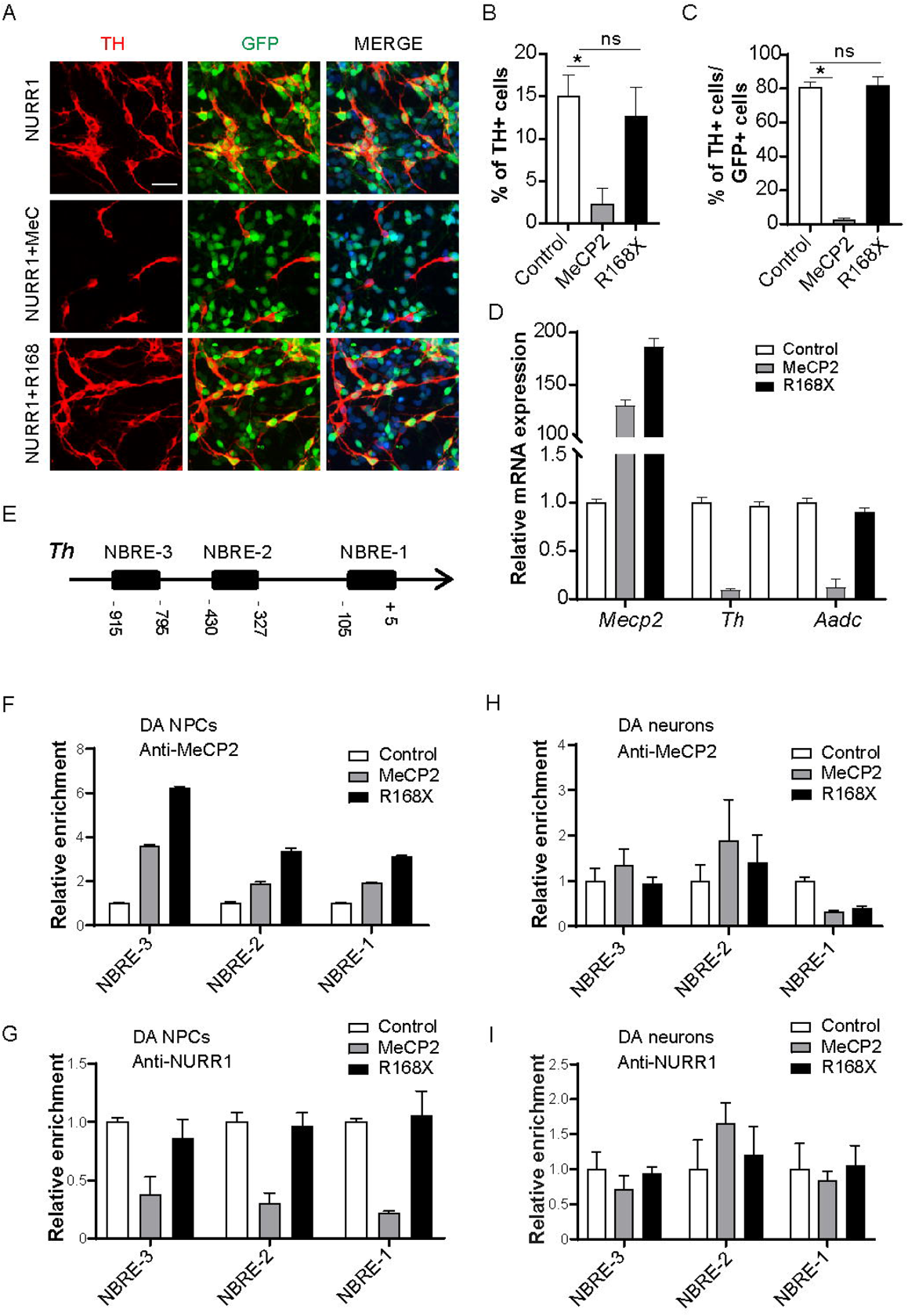
Roles of MeCP2 in NURR1-mediated transactivation of dopamine (DA) phenotype genes. (A, B) Rat cortical neural precursor cells (NPCs) were transduced with NURR1, NURR1+MeCP2 and NURR1+R168X and allowed for spontaneous differentiation for 3 days. R168X is a truncated mutation of MeCP2 which does not contain transcription repression domain. ZsGreen1 fluorescent protein (GFP) marks cells transduced with MeCP2, R168X or control vector. Representative images and quantification of cells expressing TH are shown as (A) and (B), respectively. (C) Quantification of MeCP2 or R168X-transduced cells expressing TH. (D) Transcriptional repression of NURR1-mediated expression of DA phenotype genes by MeCP2 but not R168X. (E) Schematic of 1kilo bp length of rat *Th* promoter and locations of three Nurr1 binding sites (NBREs). (F-I) Chromatin immunoprecipitation (ChIP) analyses were performed to measure the DNA binding of MeCP2 (F and H) and of NURR1 (G and I) onto NBREs in DA NPCs and DA neurons transduced with MeCP2 or R168X. For ChIP assays, one representative data from two independent experiments is shown. Data are expressed as mean ± S.D. Scale bar represents 20 μm. For immunostaining experiments, cell numbers were counted in 10 random areas of each culture coverslip. Data represent mean ± S.E.M. n=3 independent culture. * *P* < 0.05, ns, not significant; one-way ANOVA with Tukey’s post hoc analysis.

A large number of studies including ours have reported that recruitment of NURR1 onto the promoters of DA phenotype genes as the molecular mechanism for the initiation of their transcription. In particular, NURR1 transactivates *Th* gene expression through binding to NURR1 binding motif namely NBRE on *Th* promoter. Therefore, we performed a series of ChIP experiments to examine how MeCP2 interacts with NURR1 on the promoter of *Th* gene in both DA NPCs and DA neurons. Within 1 kilo bp distance upstream to transcription starting site of mouse *Th* gene, there are three NBREs (Fig. 3E). When overexpressed in DA NPCs, MeCP2 occupancy of all three NBREs was remarkably increased (Fig. 3F), whereas the binding of NURR1 was reduced (Fig. 3G). Interestingly, the binding of R168X was also increased, but it did not decrease the binding of NURR1. The best explanation of this result is that the binding of NURR1 to NBRE sites might be interfered by the TRD of MeCP2, rather than a competitive binding for NURR1 and MeCP2.

By contrast, when overexpressed in DA neurons, enrichment of NURR1 onto NBREs was not altered by overexpression of either MeCP2 or R168X (Fig. 3I). Importantly, the recruitment of MeCP2 onto NBREs was not increased by overexpression of MeCP2 or R168X (Fig. 3H), raising the possibility that the *Th* gene in DA neurons was not accessible to direct binding of MeCP2. Taken together, the data presented here demonstrate differential impact of MeCP2 overexpression on NURR1-mediated transactivation of DA phenotype genes in DA NPCs and DA neurons.

### MeCP2 competes TET1 to block DA subtype specification

We previously have revealed an essential role of TET1-mediated DNA de-methylation for NURR1 to transactivate the expression of DA phenotype genes (He et al., 2015). Therefore, we hypothesized that the DNA methylation status of DA phenotype genes distinguished the DNA binding and function of MeCP2 in DA NPCs and DA neurons. Consistent with the TET-assisted and bisulfite sequencing data in our previous study, hydroxymethylated/methylated DNA immunoprecipitation (hMeDIP/MeDIP) analyses confirmed that the 5mC level within 1 kilo bp of *Th* promoter was decreased along DA neuronal differentiation, in concert with the increase of 5hmC level (Fig. 4A). Accordingly, enrichment of TET1 on *Th* promoter, especially on the NBREs, was significantly increased (Fig. 4B), indicating that TET1 was responsible for the 5mC oxidation of *Th* promoter along DA neuronal differentiation. Interestingly, overexpression of MeCP2 in DA NPCs caused increased level of 5mC and decreased level of 5hmC after differentiation, suggesting that the natural alterations of these epigenetic marks along with differentiation were prevented by premature MeCP2 expression (Fig. 4C). In addition, ChIP analysis demonstrated that TET1 was much less recruited onto *Th* promoter by MeCP2 overexpression in DA NPCs (Fig. 4D). These results suggested that MeCP2 overexpression in DA NPCs abolished DNA de-methylation of DA phenotype genes through interfering TET1 function.

**Figure 4.**
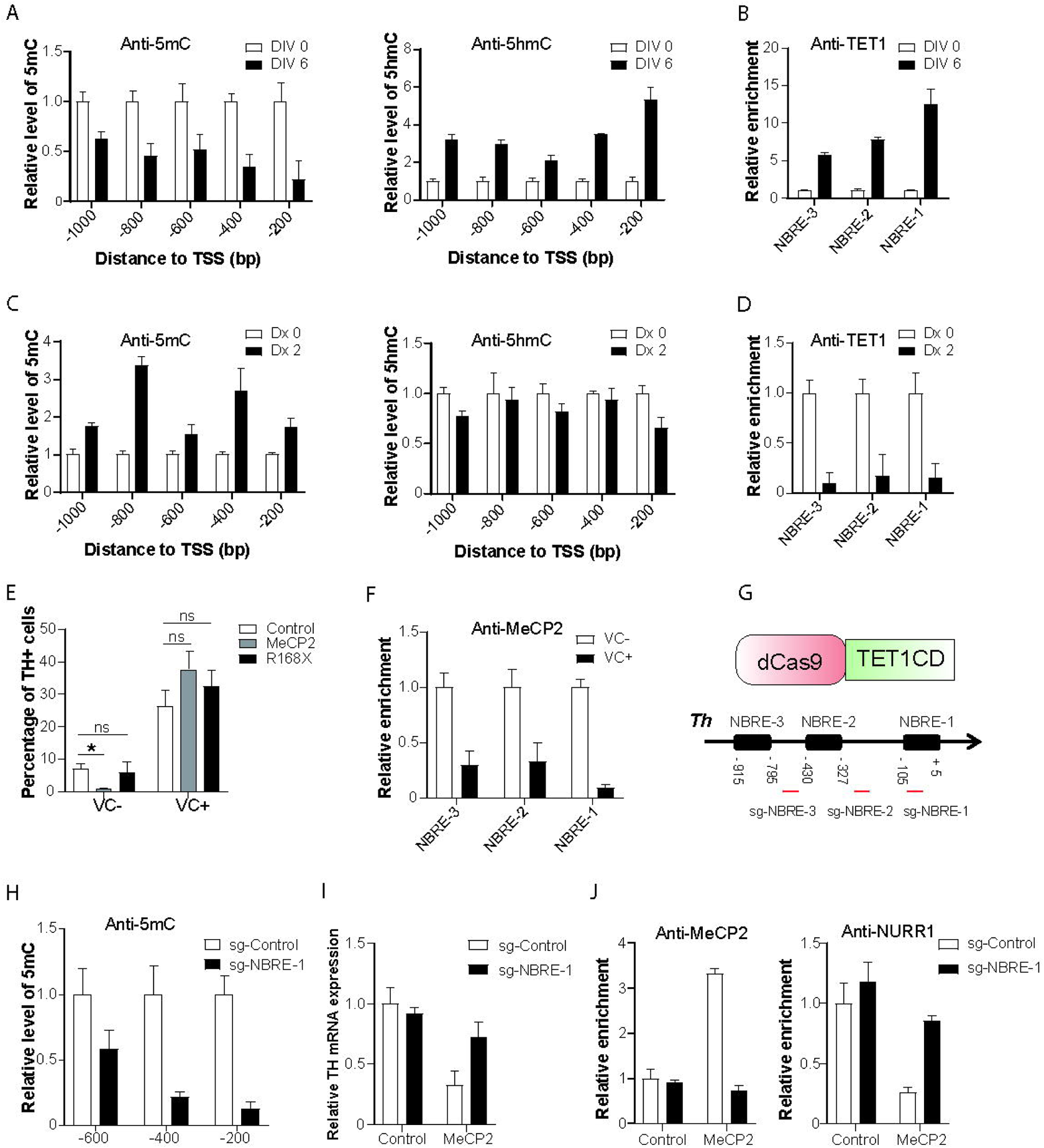
DNA de-methylation in dopamine neural precursor cells (DA NPCs) prevents MeCP2 overexpression-induced transcriptional repression of DA phenotype genes. (A) DNA methylation (5mC) and hydroxymethylation (5hmC) states within 1kilo bp promoter region of DA phenotype gene *Th* at differentiation day (DIV) 0 and DIV 6. (B) Chromatin immunoprecipitation (ChIP) experiment showing increased TET1 recruitment onto three NURR1 binding sites (NBREs) within *Th* promoter along with differentiation. (C) Doxycycline (Dx)-induced overexpression of MeCP2 in DA progenitors caused DNA methylation of *Th* promoter after DA neuronal differentiation. (D) TET1 recruitment onto NBREs within *Th* promoter at DIV 6 was reduced by overexpression of MeCP2 in DA NPCs. (E) Quantification of TH+ cells at DIV 6. Vitamin C (VC) was added in DA NPCs overexpressing MeCP2 or truncated mutation R168X from DIV 0 to DIV 6. Data represent mean ± S.E.M. n=3 independent culture. * *P* < 0.05, ns, not significant; one-way ANOVA with Tukey’s post hoc analysis. (F) Enrichment of MeCP2 onto NBREs is reduced by VC in MeCP2-overexpressed DA NPCs. (G) Schematic showing strategy of targeted recruitment of inactive Cas9-TET1 catalytic domain (dCas9-TET1CD) guided by small guiding RNAs (sgRNAs) onto three NBREs within *Th* promoter. (H) Targeted de-methylation guided sgRNA targeting the first NBRE on *Th* promoter (sg-NBRE-1) in DA NPCs caused decreased DNA methylation at and near local DNA regions after differentiation. (I) Effect of targeted de-methylation of NBRE-1 in DA NPCs with or without MeCP2 overexpression on gene expression of *Th* after differentiation. (J) ChIP experiments showing the effect of targeted de-methylation of NBRE-1 in DA NPCs with or without MeCP2 overexpression on recruitments of MeCP2 and NURR1 onto NBRE-1 after differentiation. One representative data from two independent experiments is shown. Data are expressed as mean ± S.D.

To validate whether DNA methylation of *Th* promoter is necessary for MeCP2 binding in DA NPCs, vitamin C (VC) was added to MeCP2-overexpressed DA NPCs for the purpose of a global DNA de-methylation. As previously reported (He et al., 2015), VC significantly increased the yield of TH+ cells. As shown in Fig. 2D, in the absence of VC, overexpression of MeCP2 but not R168X decreased the yield of TH+ cells. However, addition of VC completely reversed the impact of MeCP2 overexpression (Fig. 4E). Furthermore, VC remarkably decreased the enrichment of MeCP2 on the NBREs of *Th* promoter in MeCP2-overexpressed DA NPCs (Fig. 4F). These findings suggested direct associations between DNA de-methylation and *Th* gene expression with MeCP2 binding in the process of DA subtype specification. To further improve the specificity of DNA methylation, we utilized a TET1 catalytic domain (TET1CD)-mediated targeted DNA demethylation lentivirus system which has efficiently converted NPCs fate into astrocytes by targeting the methylation status of GFAP promoter(Morita et al., 2016). Single guide RNAs (sgRNAs) targeting sequences near each of three NBREs were incorporated into the vector (Fig. 4G). MeDIP experiment confirmed that sgNBRE-1 was the most potent guide RNA that was able to induce decreased levels of 5mC at and near the targeting region in DA progenitors (Fig. 4H). Co-introducing sgNBRE-1 with MeCP2 into DA NPCs attenuated the repression of *Th* expression (Fig. 4I). Furthermore, ChIP analysis demonstrated that sgNBRE-1 caused decreased MeCP2 binding and increased NURR1 binding on the first NBRE in the context of MeCP2 overexpression (Fig. 4J). Taken together, these results indicate that prematurely-expressed MeCP2 preferentially bound to hypermethylated promoters of DA phenotype genes in DA NPCs, thereby preventing DA subtype specification through interfering TET1-mediated DNA de-methylation.

### Tet1 counteracts MeCP2 to maintain DA phenotype in DA neurons

As demonstrated above, the *Th* promoter in DA neurons was hypomethylated in comparison to that in DA NPCs, and it permitted no further enrichment of MeCP2 when overexpressed in DA neurons. These results appeared to suggest an association of DNA hypomethylation with the constitutive expression of DA phenotype genes in DA neurons. Opposite to its role as a global transcriptional repressor in NPCs, recent studies have revealed MeCP2 as a global activator in differentiated neurons. An intriguing question is whether the role of MeCP2 as a transcriptional repressor of DA phenotype genes is retained during DA neuronal differentiation. To validate this, we employed a well-defined shRNA-mediated TET1 inhibition (He et al., 2015) to prevent the transition of 5mC to 5hmC in differentiating DA neurons after the DA subtype specification (from DIV4). MeDIP analysis demonstrated that TET1 knockdown caused efficient DNA methylation on *Th* promoter (Fig. 5A), suggesting that TET1 was responsible for not only the conversion but also the maintenance of DNA hydroxymethylation. Concomitantly, MeCP2 enrichment was significantly increased at NBRE-l. Overexpression of MeCP2 further exacerbated this recruitment (Fig. 5B). As a consequence, enrichment of NURR1 was reduced (Fig. 5C), and gene expression of *Th* was compromised (Fig. 5D).

**Fig 5.**
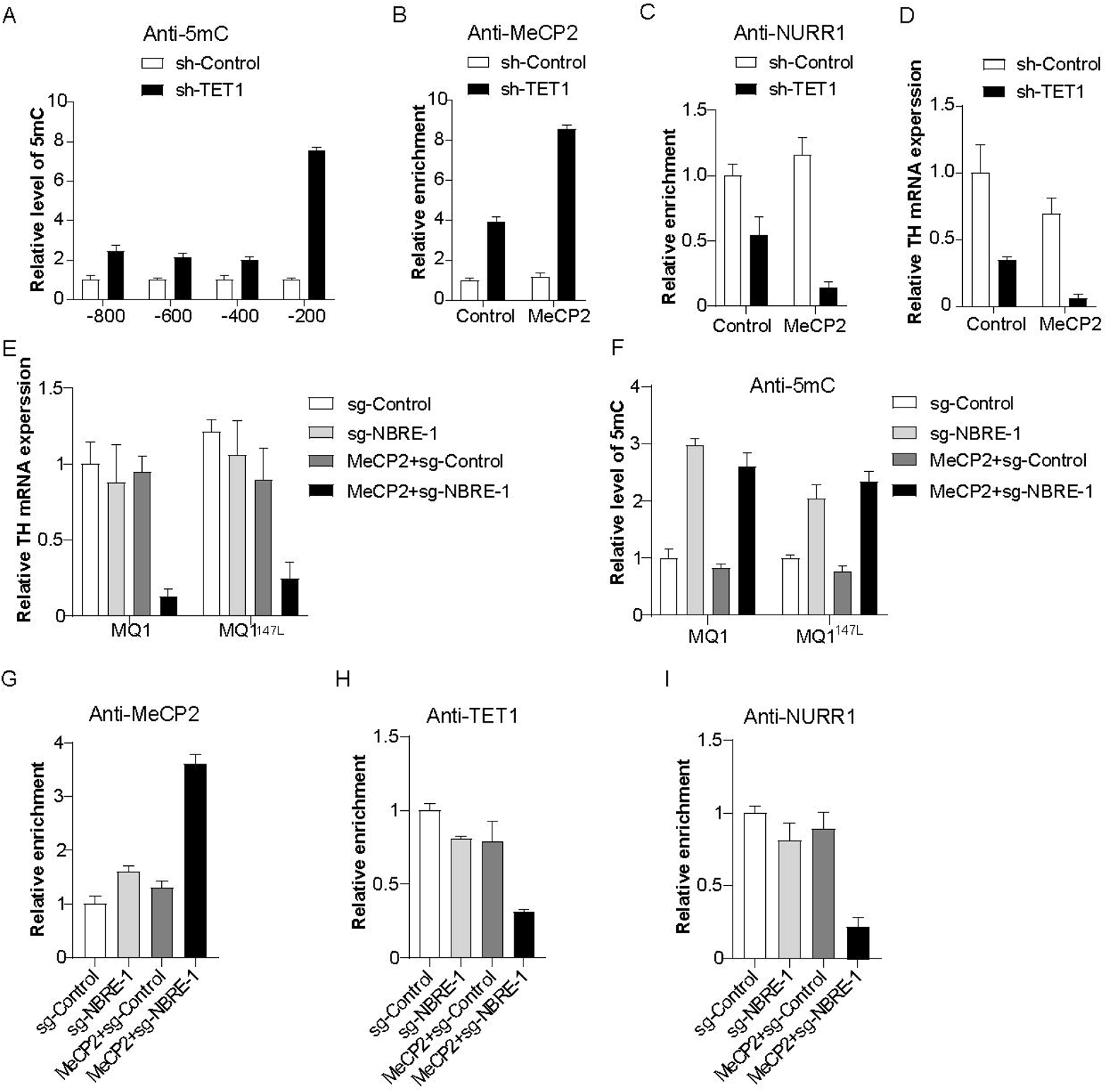
Targeted DNA methylation in dopamine (DA) neurons permits overexpressed MeCP2 counteracting TET1-dependent DA subtype maintenance. (A) Inhibition of TET1 by short hairpin RNA in DA neurons causes DNA methylation of *Th* promoter. (B and C) Effect of TET1 inhibition on recruitment of MeCP2 (B) and NURR1 (C) onto the first NURR1 binding site (NBRE-1) in DA neurons with or without MeCP2 overexpression. (D) Effect of TET1 inhibition on *Th* gene expression in DA neurons with or without MeCP2 overexpression. (E and F) Targeted recruitments of dCas9-MQ1 and its activity-enhancing mutation MQ1^147L^ onto NBRE-1 within *Th* promoter in normal or MeCP2-overexpressed DA neurons showed different impact on *Th* gene expression (E) and DNA methylation (F). (G-I) Effects of targeted DNA methylation of NBRE-1 in normal or MeCP2-ovexpressed DA neurons on recruitments of MeCP2 (G), TET1 (H) and NURR1 (I) onto NBRE-1. One representative data from two independent experiments is shown. Data are expressed as mean ± S.D.

Using a luciferase reporter assay, we previously have shown that *Th* gene expression is extremely sensitive to CpG methyltransferase MQ1-induced DNA methylation. To exclude the overall regulation effect of sh-TET1, we employed an inactive Cas9-mediated targeted DNA methylation system involving MQ1 and its mutant MQ1^147L^ with better enzyme activity (Lei et al., 2017) to manipulate targeted methylation of *Th* promoter in DA neurons. Based on targeted de-methylation results above, we focused on sgRNA targeting NBRE-1. MQ1 and MQ1^147L^ successfully caused more than 50% *Th* gene silencing in MeCP2-overexpressed DA neurons (Fig. 5E). MeDIP analysis of NBRE-1 region confirmed that 5mC was enriched by sg-NBRE-1 guided MQ1 and MQ1^147L^ (Fig. 5F). As a consequence, ChIP analysis demonstrated an increased recruitment of MeCP2 onto NBRE-1 in DA neurons experiencing overexpressed MeCP2 and MQ1-mediated targeted methylation (Fig. 5G). These results support the notion that MeCP2 functions as a transcriptional repressor that specifically recognizes methylated CpG sites in both DA NPCs and DA neurons. Conversely, decreased recruitments of TET1 and NURR1 were detected at NBRE-1 region (Fig. 5H and 5I), suggesting a counteracting role of MeCP2 with transcription activators of DA phenotype genes.

## Discussion

Here we performed a series of gain-of-function experiments to investigate the effects of abnormal timing of MeCP2 expression on midbrain DA neurogenesis, particularly on the acquisition and maintenance of DA subtype. Evidence exist for a long time in attempt to establish a direct link between MeCP2 and catecholamine system in CNS (Nomura et al., 1985). For instance, abnormal concentrations of biogenic amines have been observed in brains of MeCP2-null mouse and cerebrospinal fluid of Rett syndrome patients (Roux and Villard, 2010). However, these alterations are supposed to be independent of prenatal DA neuron development, as MeCP2-knockout embryonic stem cells are capable of differentiating into TH-expressing DA neurons in vitro (Okabe et al., 2010), and rescue of MeCP2 expression in Rett syndrome mice in postnatal stage rectified neuronal and behavior defects (Lang et al., 2013; Robinson et al., 2012). Based on these loss-of-function studies, it seems reasonable to conclude that the major role of MeCP2 is to maintain DA neuron function after neuronal differentiation. However, despite of their similar postnatal behavioral deficits, little is known about whether MeCP2 loss-of-function and gain-of-function share similar developmental pathways and molecular mechanisms. Our findings provide compelling in vitro evidence for a novel role of MeCP2 in DA neuron differentiation, proposing that controlling the timing and restricting the expression level of MeCP2 in DA NPCs is necessary for DA subtype specification. This information could be further exploited to induce efficient generation of functional DA neurons in the purpose of stem cell therapy and regenerative medicine. On the other hand, the gain-of-function of MeCP2 is genetically correlated with MeCP2 duplication or triplication syndrome. It remains to be determined whether precocious expression of MeCP2 indeed occurs in patients with these syndromes and MeCP2-overexpressing mice.

MeCP2 functions as transcriptional activator or repressor depending on genes and regulatory context (Chahrour et al., 2008). Our data support the notion that MeCP2 is a transcriptional repressor for DA phenotype genes regardless of developmental stage. This is consistent with previous studies that identified *Th* and *Aadc* as direct target genes of MeCP2 (Urdinguio et al., 2008; Yang et al., 2011). Furthermore, MBD and TRD are both indispensible for MeCP2-mediated repression, as preventing DNA binding of MeCP2 or overexpressing TRD mutation R168X failed to repress *Th* gene expression. The function of MBD determines whether or not MeCP2 is recruited to DNA sequences, allowing TET1-mediated de-methylation to distinguish DNA binding of MeCP2 in DA NPCs and DA neurons. The function of TRD is to recruit other repressive complexes, so as to block the recruitment of other activators such as NURR1 and TET1. Interestingly, one study using MeCP2-overexpressing mice demonstrate that the duplication toxicity requires both domains to be functional (Heckman et al., 2014), raising the possibility that too much MeCP2 protein might function as a universal transcriptional repressor rather than activator.

The molecular mechanisms proposed here raise at least two neuropathological concerns (Fig. 6). First is the premature MeCP2 expression in DA NPCs. We previously have shown that the promoter regions of other DA phenotype genes such as *Aadc* and *Dat* were also hypermethylated like *Th* in DA NPCs (He et al., 2015), and the expression of these genes are also under transactivation of NURR1 (van Heesbeen et al., 2013). It is possible that the mode of action of premature MeCP2 expression proposed here is applicable to other DA phenotype genes, affecting a wider range of DA neuronal functions including DA synthesis, transport and re-uptake. Second is failed de-methylation of DA phenotype genes in DA neurons. In Figure 5, MeCP2 overexpression in DA neurons enhanced the gene repression effect induced by targeted methylation of *Th* promoter, suggesting that the maintenance of DA phenotype genes in DA neurons is sensitive to DNA methylation. As MeCP2 is highly expressed in adult midbrain DA neurons, genetic deficiency such as TET1 mutation or environmental cues that induce DNA methylation might work as potential risk factors for the proper functioning of midbrain DA neurons. However, although DNA methylation was induced in both MeCP2-overxpressed and normal DA neurons, the transcription of *Th* gene was only repressed in MeCP2-overexpressed DA neurons. Consistently, DNA bindings of repressors and activators were also not altered in normal DA neurons experiencing MQ1-mediated targeted DNA methylation. These results might suggest the existence of an unidentified regulation mode that protects DA neurons from DNA methylation-related gene silencing.

**Fig 6.**
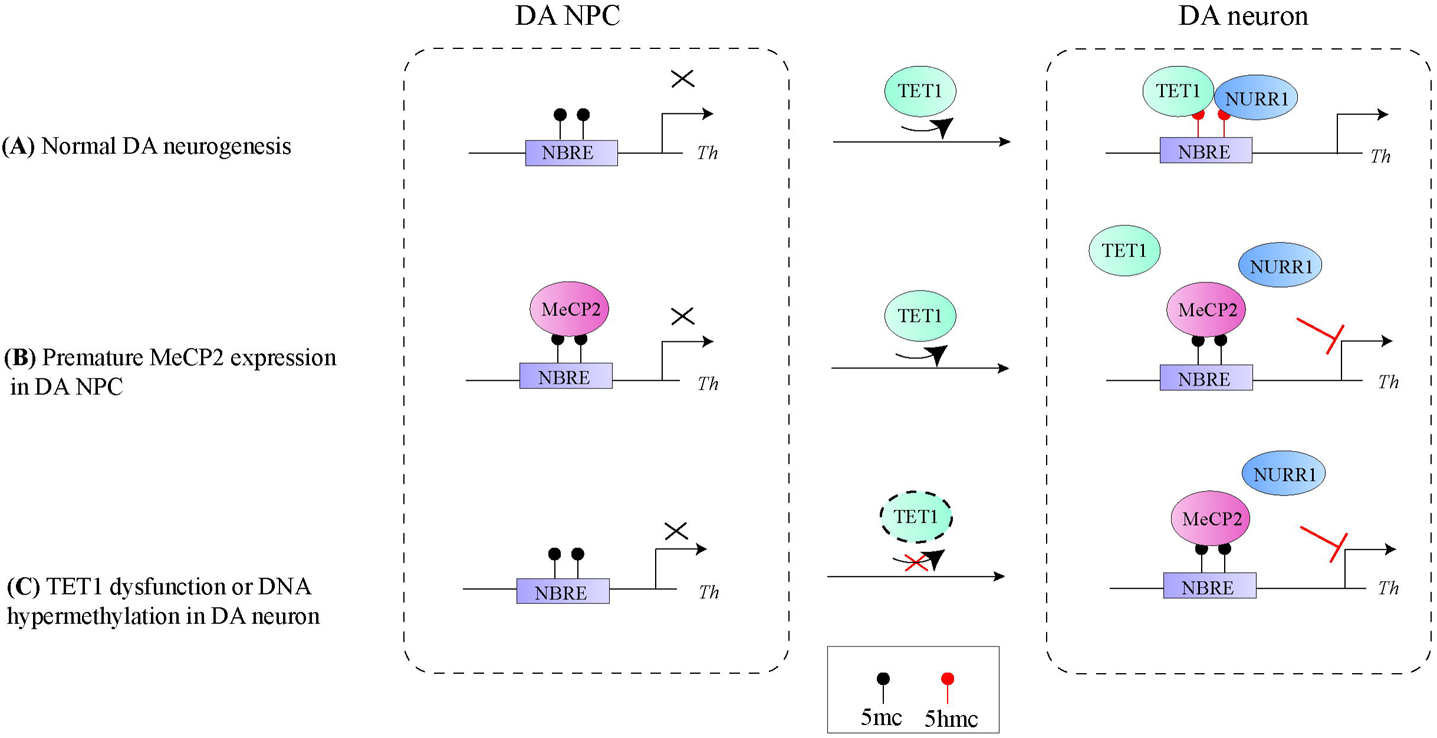
Proposed model illustrating mechanisms underlying differential effects of timing and dose of MeCP2 expression on dopamine (DA) subtype specification and maintenance during midbrain DA neurogenesis. *Th* serves as a representative for DA phenotype genes. (A) Under normal development, DA subtype is not yet specified in DA neural precursor cells (NPCs) due to hypermethylation of the promoters of DA phenotype genes and lack of NURR1. Along with differentiation, TET1 induces gene-specific DNA de-methylation, in particular of NURR1 binding sites (NBREs), which allows recruitment of NURR1 for gene transactivation. The expression of DA phenotype genes is maintained after differentiation in DA neurons. (B) MeCP2 is normally not expressed in DA NPCs before subtype specification. When MeCP2 is prematurely expressed, for instance, in MeCP2 duplication syndrome, the hypermethylated promoters of DA phenotype genes become targets for MeCP2 recruitment and its mediated gene repression. This prevents TET1-mediated DNA de-methylation of DA phenotype genes along with DA neuronal differentiation. As a consequence, DA NPCs and neurons after differentiation are not able to acquire DA subtype. (C) In DA neurons, TET1 continues de-methylating DA phenotype genes, allowing constitutive NURR1 binding and maintenance of DA phenotype. TET1 dysfunction leads to hypermethylation of DA phenotype genes, allowing binding of excess MeCP2, failed NURR1 binding and subsequent loss of DA subtype.

## Materials and Methods

### Cell culture

DA NPCs were extracted from ventral regions of mouse E10.5 mesencephalon, mechanically dissociated into single cells and plated onto poly-L-ornithine and fibronectin double-coated glass coverslips and culture plates. Cells were allowed for proliferation by daily addition of basic fibroblast growth factor and epidermal growth factor in Neurobasal media supplemented with N2 and B27 supplement. DA neuron differentiation was conducted by withdrawal of mitogens from culture media for 3 to 6 days. For NURR1 overexpression experiments, NPCs were extracted from rat embryonic day 13 cortices and cultured by similar approach to mouse DA progenitors, except that only basic fibroblast growth factor was used for proliferation and B27 supplement was not included in the media. All culture media were changed every other day and cells were maintained at 37℃ in humidified 5% CO2 incubators.

### Plasmids and gene delivery

cDNAs encoding full length mouse MeCP2 and NURR1 was generated by reverse transcriptase-PCR from RNA extracted from adult mouse brain. Plasmid expressing Dx-inducible MeCP2 was made by cloning the cDNA into FUW-TetO-MCS vector (Addgene plasmid #84008). Plasmid expressing MeCP2 and GFP tagging was made by cloning the cDNA into pLVX-EF1α-IRES-ZsGreen1 vector (TaKaRa plasmid #631982). R168X mutation was introduced by site-directed mutagenesis. Plasmid expressing NURR1 was made by cloning the cDNA into pLVX-based vector. Plasmid expressing rat-specific shTET1 was purchased from Origene (#309902). For targeted DNA de-methylation, sgRNAs were cloned into pPlatTET-gRNA2 vector (Addgene plasmid #82559). Plasmids expressing sgRNAs were made by cloning the annealed oligo sgRNA sequences into PX459-based vector (Addgene plasmid #62988). The oligo sequences for sg-NBRE-1, -2, -3 are 5’-GAGGGGCTTTGACGTCAGCC-3’, 5’-AATTAGATCTAATGGGACGG-3’,5’-ACTTTGTTACATGGGCTGGG-3’, respectively.

For the production of viruses, the virus vectors and 2nd generation packaging vectors were introduced into HEK293T packaging cell line by transient transfection with Lipofectamine 2000 (Invitrogen). One day later, supernatants were harvested, concentrated with PEG8000 (Sigma), and stored until use. For viral transduction, cells were incubated with the viral supernatant containing polybrene (4 μg/ml, Sigma) for 6 hours.

For targeted DNA methylation, MQ1 and MQ1^Q147L^ plasmids (Addgene plasmid #89634, 89637) and sgRNAs were co-transfected into DA neurons with Lipofectamine Stem (Invitrogen) for 6 hours.

### Immunofluorescent staining and imaging

Cells were fixed with 4% paraformaldehyde for 20 minutes, permeabilized and blocked with phosphate buffered saline with 0.3% Triton-X100 and 1% bovine serum albumin for 40 minutes, then incubated with first antibodies diluted with blocking solution at 4℃ overnight. Alexa Fluo series of second antibodies (Thermo Scientific) were applied accordingly for two hours at room temperature. Cells were finally mounted in 4’,6-diamidino-2-phenylindole (DAPI) and examined using fluorescence microscope (Leica DMi8). Embryonic and postnatal ventral midbrain tissues were fixed with 4% paraformaldehyde and dehydrated with 30% sucrose overnight, and cryosectioned at 14 μm thickness. The first antibodies used include rabbit anti-MeCP2 (Cell Signaling Technology), mouse anti-NURR1 (R&D Systems), mouse anti-TH (Sigma), rabbit anti-NURR1 (Santa Cruz Biotechnology).

### Real-time PCR analysis

RNA was extracted, reverse-transcribed (TaKaRa), amplified, and applied to real-time PCR analyses (Roche). The comparative cycle threshold method was used for quantification. Each experiment was repeated at least once to guarantee same trend of gene expression. Data from one experiment was used to generate representative histogram. The primer sequences are listed in Supplemental Table 1.

### ChIP, MeDIP and hMeDIP

For ChIP experiments, cells were cross-linked with 1% paraformaldehyde for 15 minutes and Chromatins were sheared into an average 200–400 bp in length by sonication (Diagenode) and immunoprecipitated with following antibodies: rabbit anti-MeCP2 (Cell Signaling Technology), rabbit anti-NURR1 (Santa Cruz Biotechnology) and rabbit anti-TET1 (Abcam). Immunoprecipitated DNA fragments were collected by magnetic beads (Active Motif), purified, and subjected to real-time PCR using primers specific to regions spanning three NBREs on *Th* promoter. Data were normalized to values of the input DNA. For MeDIP and hMeDIP experiments, genomic DNA were extracted from cells, sheared, immunoprecipitated, collected and subjected to real-time PCR in a similar way as ChIP with following antibodies: mouse anti-5mC (Abcam) and rabbit anti-5hmC (Active Motif). The primers are listed in Supplemental Table 1.

### Cell counting and statistics

Immunoreactive or DAPI-stained cells were counted in at least 10 random regions of each culture coverslip using an eyepiece grid at a magnification of 50 to 400X. Data are expressed as mean ± S.E.M. of three to five independent cultures. Statistical comparisons were made using Student’s t-test or one-way ANOVA with Tukey’s post hoc analysis (Graphpad Prism).

## Acknowledgments

This work was supported by the National Natural Science Foundation of China [31701287 to X-B.H.].

## Author Contributions

X-B.H. conceived the study and wrote the manuscript. X-B.H. performed cell culture, immunostaining and epigenetic experiments. F.G. performed molecular cloning, virus production, PCR experiments and helped with data collection and data analysis. K.L. and J.Y. were involved in data analysis and figure arrangement. S-H.L. provided technique support and supervised epigenetic experiments. All authors have read and approved the final version of the manuscript.

