## Supplemental Table S1 for "Premature MeCP2 Expression Disturbs the Subtype Specification of Midbrain Dopamine Neurons"

**Supplemental Table S1.** Oligonucleotide primers

| Primer name | Forward | Reverse |
| --- | --- | --- |
| Primers for ChIP analysis | | |
| Th NBRE-1 | AGAGGATGCGCAGGAGGTAGGAG | GTCCCGAGTTCTGTCTCCAC |
| Th NBRE-2 | TCCTGGAGGGGACTTTATGA | CTGGATTTCCTAAGGGCTCA |
| Th NBRE-3 | GGGTGTGGATGCTAACTGGA | AGTGGTAGCCCCATTCTCAG |
| Primers for MeDIP and hMeDIP analysis | | |
| Th_1 (-200,0) | GCGACAGTGGATGCAATTAG | CCTCTTAAAGGCCAGGCTGA |
| Th_2 (-400,-200) | GGGGACTTGAAGACATCCAA | CCCTAGACACGACATGAAGACA |
| Th_3 (-600,-400) | TAGGGAGATGCCAAAGGCTA | AGTGTATGTGCTGGCACTGG |
| Th_4 (-800,-600) | GCTCAGCATAAGTCCCCTGT | GTAAGGGCGCACTCAGTGAT |
| Th_5 (-1000,-800) | ACTCAGTGTGCTGGGTCCTC | AGGCAATGGTTCAGGACAAC |
| Primers for gene expression analysis | | |
| Th | AAGGGCCTCTATGCTACCCA | TGCATTGAAACACGCGGAAG |
| Aadc | CCTACTGGCTGCTCGGACTAA | GCGTACCAGTGACTCAAACTC |
| Mecp2 | AAGAGGGCAAACATGAACCACT | TTGCCTGCCTCTGCTGG |
| Gapdh | CTCATGACCACAGTCCATGC | TTCAGCTCTGGGATGACCTT |
